# Value-based decision-making between affective memories

**DOI:** 10.1101/2021.06.14.448275

**Authors:** Erdem Pulcu, Calum Guinea, Hannah Clemens, Catherine J Harmer, Susannah E Murphy

## Abstract

Affective biases can influence how past events are recalled from memory. However, the mechanisms underlying how discrete affective events shape memory formation and subsequent recall are not well understood. Further understanding this is important given the central role of negative biases in affective memory recall in depression and antidepressant drug action. In order to capture cognitive processes associated with affective memory formation and recall, we studied value-based decision-making between affective memories in two within-subject experiments (n=45 and n=74). Our findings suggest that discrete affective events, created by large magnitude Wheel of Fortune (WoF) outcomes, influence affective memory formation processes during reinforcement-learning (RL). After 24 hours, we show that healthy volunteers display stable preferences during value-based recall of affective memories in a binary decision-making task. Computational modelling of these preferences demonstrated a positive bias during value-based recall, induced by previously winning in the WoF. We further showed that value-based decision-making between affective memories engages the pupil-linked central arousal systems, leading to pupil constriction prior to, and differential pupil dilation after the decision onset depending on the valence of the chosen options. Taken together, we demonstrate that mechanisms underlying human affective memory systems can be described by RL and probability weighting models. This approach could be used as a translational assay to study the effects of novel antidepressants.

## Introduction

Human social life is arguably the most complex in the animal kingdom, enriched by our ability to express and infer from others a wide spectrum of emotions. The breadth of this affective repertoire, along with our tendency to process positive and negative information asymmetrically^1,2^, makes us prone to affective biases that can shape not only our present experiences, but also how we recall events from the past. For example, the study of eyewitness memory has highlighted that centrally relevant details (e.g. the characteristics of a criminal or the weapon used in a crime) from emotional events are remembered more accurately than non-affective content^3^. It is known that humans exhibit an asymmetry in affective information processing (hence forth “affective bias”). In healthy volunteers, affective bias is more frequently observed in favour of positive events, and spans across multiple domains including perception, attention, reinforcement learning (RL), and memory ^2,4,5^. In psychiatric conditions such as major depressive disorder (MDD), negative affective biases (i.e. preferential processing of negative relative to positive information)^6–10^ have been shown to play a role in the development and maintenance of symptoms^11–13^. Nevertheless, mechanisms underlying how discrete affective events induce biases that can influence learning and subsequent memory recall in humans remain elusive.

Recent preclinical work has further elucidated how discrete affective events can influence memory-guided value-based decisions, and how these can be targeted pharmacologically. Stuart, et al. ^14^ (2015) demonstrated that ketamine, a non-competitive N-methyl-D-aspartate (NMDA) receptor antagonist known to have rapid antidepressant (AD) effects^15–17^, injected into mouse medial prefrontal cortex (mPFC), attenuates negative memory biases. This effect was shown in a decision-making assay in which rodents were probed to choose between two substrates with equal nutritional value: one previously paired with an anxiogenic compound (FG7142) and another paired with saline during learning. This finding demonstrates the malleability/plasticity of cognitive processes underlying negative affective biases and has important implications for understanding the mechanisms of rapid AD treatment of depression. Translating preclinical paradigms which are developed within the constraints of animal models of disease is critically important for unifying human and animal work under a single mechanistic umbrella^18,19^. This interdisciplinary approach can help to speed up drug discovery in psychiatry^20^. In the current work, we will describe a behavioural assay founded in RL and value-based decision-making, translating the essence of the rodent assay from Stuart et al., (2015) to shed light on cognitive mechanisms underlying value-based recall of affective memories in humans.

In humans, there is evidence demonstrating that discrete affective events influence subsequent value-guided choice. Using a Wheel of Fortune (WoF) manipulation, Eldar and Niv ^21^ (2015) provided quantitative evidence showing that individuals who scored highly on a mood instability measure and won money in the WoF draw preferred probabilistic slot machines they experienced immediately after the draw, whereas those who lost in the WoF draw preferred those slot machines that preceded the draw, even though expected values of the slot machines on either side of the WoF draw were comparable. In the current work, we adopted a similar experimental design to manipulate participants’ affective state. Unlike Eldar and Niv (2015), who assessed the impact of such affective events on participants’ value-based decisions shortly after the WoF manipulation, we tested participants’ preferences between abstract information learnt through RL, and up to 4 days later. Thus, in our work, preference biases observed in participant choice behaviour would be driven by “affective memories” based on information encoded through RL in earlier stages of the experiment (see Methods for further details). This within-subject approach captures the essence of the rodent assay and it is also similar to the methodology used in a recent study which investigated serotonergic modulation of learning and memory-based decision-making processes^22^. Here, use of the RL framework also ties in with the importance of implementing computational methods for understanding the mechanisms underlying affective biases. This is important because recent RL studies demonstrated that negative affective biases, which are known to be causally linked to symptoms of depression^23^, may develop even in healthy volunteers as a rational response to environmental contingencies^24^ and relate to poor filtering of informative negative experiences from uninformative ones^25^. In these previous studies, we demonstrated that the information content of negative affective events engages the pupil-linked central arousal systems ^24^. In the current study, we used pupillometry to expand on these previous findings and to investigate whether value-based recall of affective memories also engages central arousal systems.

The aim of the current study was to test whether experimentally induced changes in emotional state influence human choice behaviour during RL **(Figure 1)**. Secondly, we investigated whether nonclinical volunteers display a positive bias during value-based decision-making between affective memories. Finally, we investigated whether this process engages the pupil-linked central arousal systems. We predicted that discrete affective events should have a significant and differential influence on human RL. We predicted that a non-clinical population would overall display a positive bias, indicated by a preference for shapes encoded after winning on the WoF. We analysed participant choice behaviour with a well-established computational model of value-guided choice, which posits that choice preferences can be expressed in terms of weighted probabilities^26^. Finally, using a model-based analysis of pupillary data, we tested the prediction that subjective values which guide value-based decision making between affective memories will significantly influence pupil dilation.

**Figure 1.**
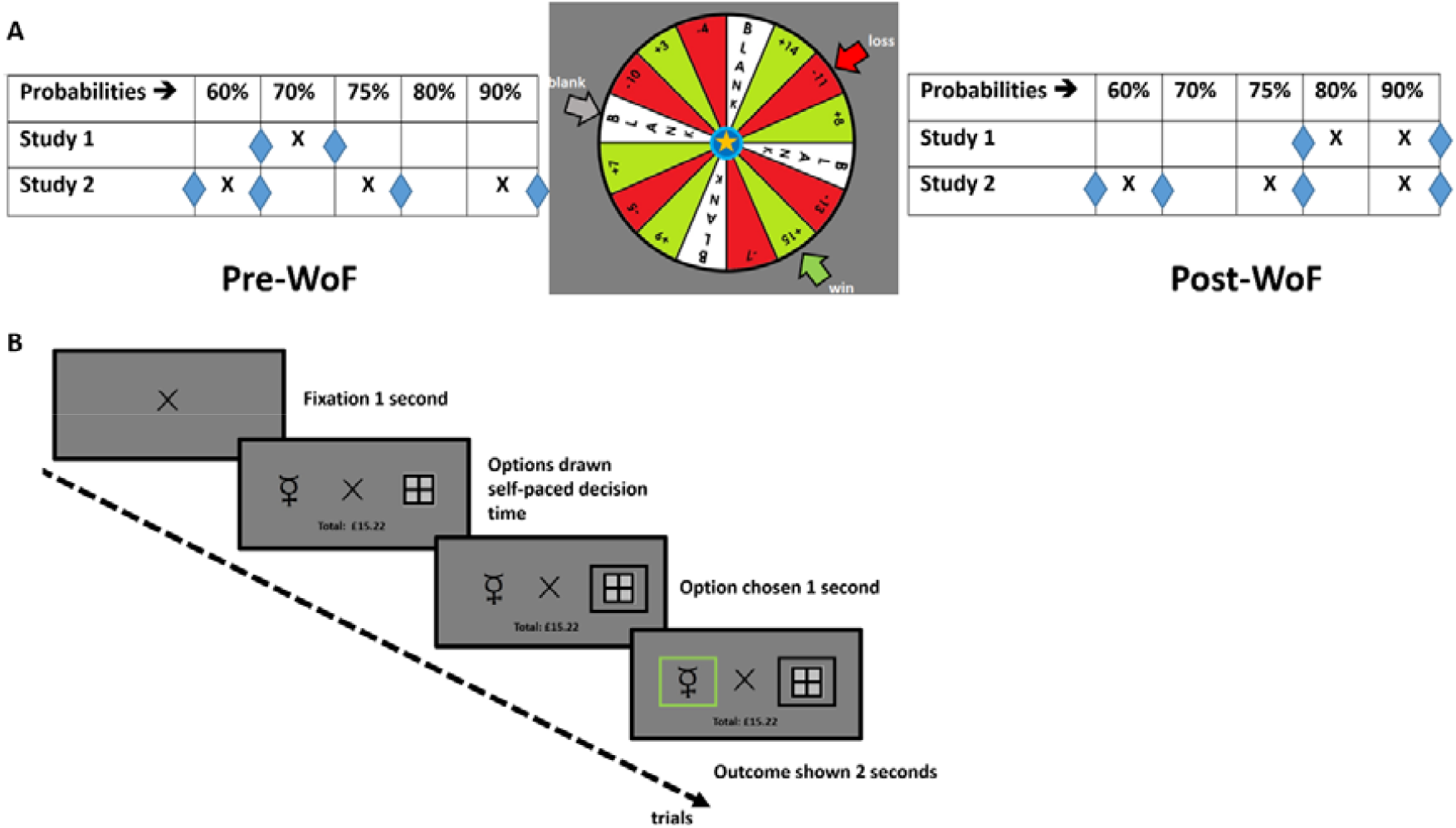
(A) An overview of the studies. Two studies were conducted to understand how discrete affective events influence human reinforcement learning. In Study 1 (conducted in the laboratory), participants completed the task on three consecutive days. On each day they experienced a different WoF outcome, the order of which was counterbalanced across participants; win (£15), loss (-£11) or a blank (see middle of panel A). Participants completed a single baseline block of learning trials pre-WoF (70% reward probability) and two subsequent blocks post-WoF (90% and 80% reward probabilities). In Study 2 (conducted online) participants completed the task on two consecutive days. On each day they experienced a different WoF outcome, the order of which was counterbalanced across participants: win (£14) and loss (-£7). On each day they completed 3 blocks of learning trials pre-and-post WoF, with matched reward probabilities were matched. The table shows the probability (p) associated with the higher reward probability shape, where the probability associated with the other shape is 1-p. Blue diamond markers indicate the timepoints of happiness rating assessments. **(B) Reinforcement learning task**. After a fixation period of 1 second, participants had to choose, using the left and right arrow keys, between two abstract shapes. They were asked to choose the shape that was most likely to be rewarded (i.e. the shape associated with a higher reward probability). After participants made their choice, a black frame appeared around the chosen shape. If the choice was correct, the black frame would turn green. If the choice was incorrect, a green frame would appear around the unchosen shape. On each trial, one of the shapes was linked to a ‘win’ outcome (+2 pence) and the other shape would result in no monetary gain. The win and null outcomes were dependent on each other (probabilities add up to 1). Using trial and error, participants could infer the reward probability associated with each shape. This information could then be used to maximise their monetary reward. Participants started with £15 and their running total, displayed below the fixation cross for the duration of every trial, updated by 2p for each correct choice made. Incorrect choices did not have any monetary effect on participants’ running total.

## Materials and Methods

### Participants

Forty-five (Study 1) and seventy-four (Study 2) English-speaking healthy participants were recruited from the general public using online and print advertisements around Oxfordshire, UK. All of the participants had normal or corrected to normal vision and did not report a present or past psychiatric diagnosis, nor any serious medical condition that could impact their study participation. Participants were excluded if they were currently using psychotropic medication. Participants received monetary reimbursement for their time (£50) plus additional payment depending on their task performance across the learning and decision-making components of the experiment (£33.26-£38.40, mean±SD £37.25±0.90). The study was approved by the University of Oxford Central Ethics Committee (CUREC; ethics approval reference: R66705/RE001). All participants completed an informed consent form conforming to the Declaration of Helsinki.

### General Experimental Procedures

In Study 1, testing sessions took place over 5 consecutive days at the University of Oxford, Department of Psychiatry at Warneford Hospital. In the first visit, the participants were taken through a screening interview to assess their eligibility. Then, the participants responded to a set of demographic questions and completed a battery of psychological questionnaires. After the screening interview, the eligible participants continued with the first day of learning and completed 3 blocks of a simple RL task in order to learn the associations between shapes and rewards. In line with the aims of the study, participants’ affective state was manipulated using a WoF paradigm adapted from Eldar and Niv (2015). On each day participants experienced a different WoF outcome: win (£15), loss (-£11) or a blank (see **Figure 1** and legends about a detailed description of the experiment). We used these large magnitude WoF outcomes to experimentally induce negative or positive memory biases. To probe value-based recall of affective memories, after the training days, we asked participants to make decisions in a two-option forced-choice (TOFC) preference task in which various combinations of the abstract shapes they had learned about were paired with each other (i.e. on the last 2 days of the lab-based study, and the last day of the online study). Although no explicit feedback was given to participants in the preference test, they continued to accumulate money based on the reward probability of the chosen shape (i.e. 90% chance of winning 2p if the participant selects the shape associated with 90% reward probability in the learning phase). We were particularly interested in the pairs of shapes that had objectively identical reward probabilities but appeared after different WoF outcomes, thus should be encoded under different affective influence (**Figure 1**). Therefore, the majority of trials presented during the preference tests compared shapes of equal reward probability but under different affective influence (**Supplementary Figure 1**). All tasks were presented on a laptop running MATLAB (MathWorks Inc) with Psychtoolbox (v3.1).

In Study 2, testing sessions took place over 3 consecutive days and were delivered using an online platform (due to the global COVID-19 pandemic). We manipulated the reward probabilities in each RL block pre-and-post WoF in a balanced manner in order to investigate how discrete affective events influence human RL. Further details of experimental procedures and statistical analysis approach and computational modelling is in Supplementary Methods and Materials.

## Results

### Participants and demographics

Demographics and a summary of psychological questionnaire measures are given in **Table 1**. In both Study 1 and Study 2, depression and trait anxiety scores were highly significantly correlated (r = .57 and r = .70, both p< .001).

**Table 1.** Participant Demographics

| Measure | Study 1 (n = 45) | Study 2 (n = 74) |
| --- | --- | --- |
| | Mean ± SD | Mean ± SD |
| Age | 29.33 ± 7.98 | 33.94 ± 7.95 |
| Gender, female | 31 (69%) | 62 (84%) |
| Years of education | 16.53 ± 3.00 | N/A |
| Trait-STAI | 32.51 ± 9.11 | 32.76 ± 10.94 |
| State-STAI | 29.31 ± 7.39 | 29.83 ± 11.1 |
| BDI | 3.56 ± 4.00 | 5.15 ± 5.46 |
| BAS Drive | 6.64 ± 2.31 | 8.2 ± 2.75 |
| BAS Fun | 7.16 ± 2.01 | 8.45 ± 2.71 |
| BAS Reward | 12.02 ± 1.71 | 13.05 ± 2.59 |
| BIS | 15.69 ± 3.41 | 16.15 ± 3.92 |
| MDQ | 3.02 ± 3.65 | 4.08 ± 3.68 |
| PANAS positive affect | 34.64 ± 6.97 | 31.54 ± 10.4 |
| PANAS negative affect | 17.67 ± 12.40 | 12.21 ± 4.51 |

Trait STAI, Spielberger State-Trait Anxiety Inventory, trait form; State STAI, Spielberger State-Trait Anxiety Inventory, state form; BDI, Beck Depression Inventory; BAS, Behavioural Activation; BIS, Behavioural Inhibition; MDQ, Mood Disorder Questionnaire; PANAS, Positive and Negative Affect Schedule.

### Discrete affective events influence happiness ratings and human reinforcement learning

To test whether the WoF manipulation influenced participants’ happiness, we compared their happiness ratings immediately before (pre-WoF) and immediately after (post-WoF) the draw. Overall, participants’ ratings indicated that they felt significantly happier immediately after winning in the WoF, and felt significantly less happy immediately after losing (statistical details are available in Supplementary Results and **Supplementary Figure 2A**). Therefore the wheel of fortune was effective at modulating mood in the expected direction.

**Figure 2.**
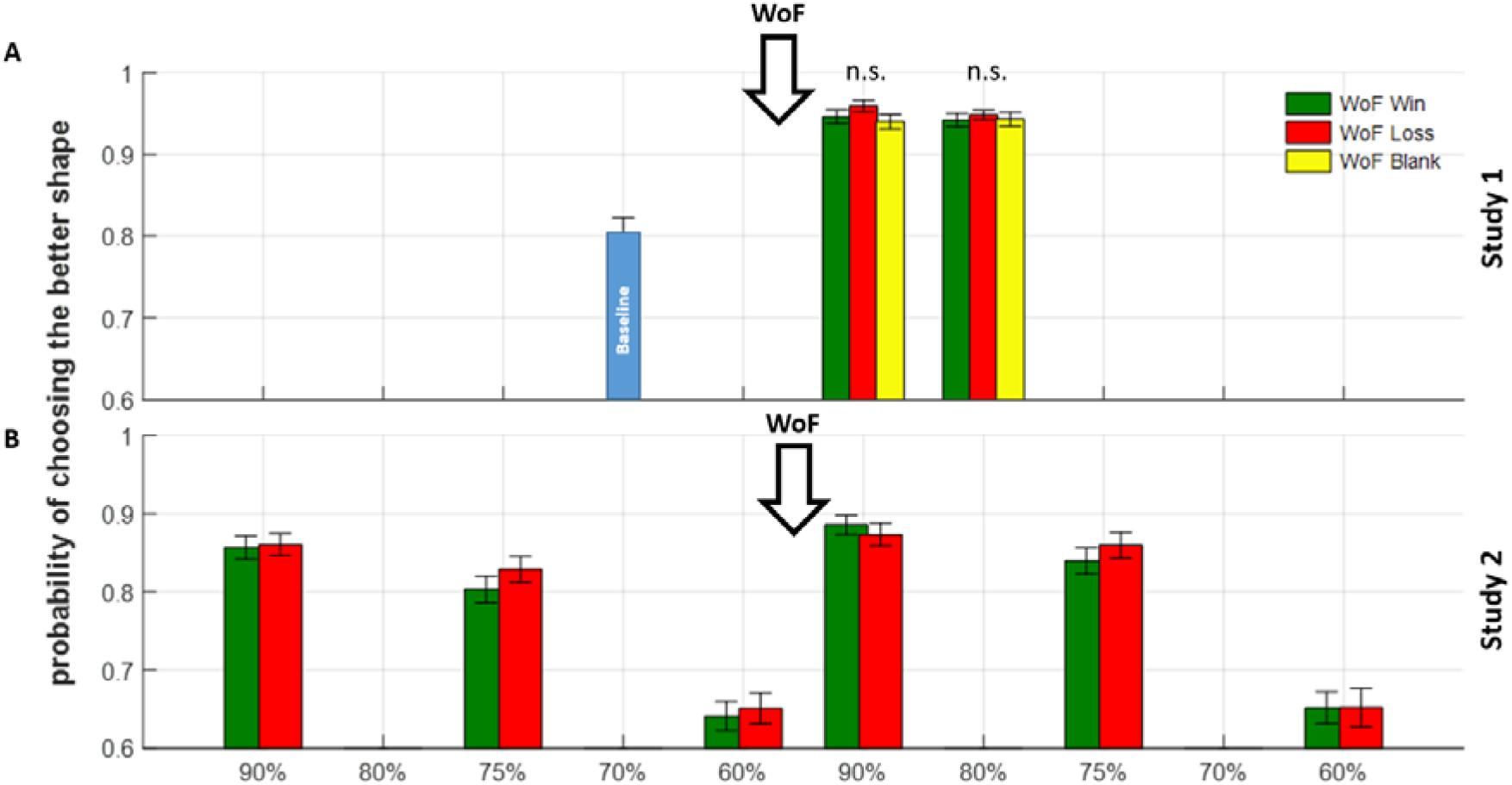
Summaries of probability of choosing the higher probability shape. **(A)** In Study 1, there were no significant differences in participant learning behaviour after the WoF, irrespective of the valence of WoF outcome. The single blue bar at 70% before the WoF shows the baseline condition. **(B)** In Study 2, we were able to compare pre-and-post-WoF learning behaviour. A repeated measures ANOVA model indicated a significant main effect of WoF influencing participant choice behaviour, reflecting an increased probability of choosing the shape associated with a high probability of reward post-WoF, irrespective of valence of the WoF outcome. Downward arrow with WoF indicates the point in which participants experienced the WoF draw within the course of their daily learning sessions. Note that, Study 2 only had win and loss outcomes in the WoF. In both panels, error bars reflect ±1 SEM. Probabilities on x-axis reflect reward probabilities from Figure 1A.

In Study 1, we were able to investigate whether the valence of discrete affective events (i.e. winning, losing or a blank outcome on the WoF) influence human reinforcement learning. A rmANOVA did not reveal any significant main effect of WoF outcome on the probability of choosing the better shape (i.e. the shape associated with higher reward probability) post-WoF. This is illustrated in **Figure 2A**, where we show that valence of the WoF outcome does not influence learning in the post-WoF blocks. Moreover, the interaction term (WoF outcome by reward probability) was not significant and there was no main effect of WoF outcome order (all p > .136). However, in this study (Study 1) we were only able to compare learning behaviour in the post-WoF blocks which were identical in terms of their reward probabilities, but we were not able to understand how learning behaviour might have changed from the pre-WoF baseline, as the reward probability in the pre-WoF block was different (i.e. 70%). We addressed this question by improving on the experimental design in Study 2 in which participants completed an identical number of blocks pre-and post-WoF with identical reward probabilities (**Figure 1A**). Due to lack of a significant main effect of WoF on learning behaviour in the blocks subsequent to it, we did not further analyse the data from Study 1 with computational models.

In Study 2, a rmANOVA (2 valence x 3 probability levels x 2 phases (i.e. pre versus post WoF RL blocks), also including the win/loss training order as a between-subjects factor) revealed, consistent with Study 1, that there was no significant main effect of valence (F(1,65) = 2.439, p = .123, **Figure 2B**), suggesting that the outcome of the WoF did not affect subsequent learning. There was, however, a significant main effect of phase (F(1,65) = 17.423, p<.001), reflecting an increased probability of participants selecting the shape associated with a high probability of reward post-WoF. There were no significant interactions and no main effect of WoF order (F(1,65) = 1.374, p = .245). In order to understand how discrete affective events influence human reinforcement learning, we further analysed participant choice behaviour in the online study using computational modelling (in Supplementary Results).

### Value-based decision-making between affective memories reveals stable preferences

In Study 1 we evaluated these preferences on two subsequent days to be able to establish the stability of value-based decision-making between affective memories (2×4 rmANOVA: 2 preference days, 4 shape valence). There was no main effect of test day on participant choice behaviour (i.e. preference day 1 versus day 2, F(1,39 = .303, p = .585), indicating that value-based decision-making between affective memories remained stable (test-retest reliability coefficient 0.883). We observed a significant main effect of shape valence (F(3,117) = 5.912, p = .001). Shapes which were learnt following a WoF win or loss were selected more frequently than shapes learnt during the baseline (pre-WoF) block and those learnt following a neutral (blank) WoF (**Supplementary Figure 6A**). Specifically, loss shapes were not preferred significantly over win shapes (day 1: (t(86)=1.1902, p=.24; day 2: t(86)=1.867, p=.07), but preferred significantly over blank shapes (day 1: t(86) = 2.857, p = .005; day 2: t(86) = 2.267, p = .026) and over baseline shapes (day 1: t(86) = 4.617, p < .001; day 2: t(86) = 4.313, p < .001), while win shapes were chosen over blank shapes (day 1: t(86) = 1.689, p = .09; day 2: t(86) = 0.465, p = .643) and over baseline shapes (day 1: t(86) = 3.435, p < .001; day 2: t(86) = 2.418, p < .018). The comparison between blank vs. baseline shapes was not significant (day 1: t(86) = 1.644, p = .10; day 2: t(86) = 1.784, p = .078). There was no significant main effect of WoF outcome order on participant choice behaviour (F(5,39) = .364, p = .870). Pairwise comparisons between equal value shape pairs are summarised in **Supplementary Table 1**.

We further investigated preferences between equal value shapes in Study 2. We observed that discrete affective events of comparable magnitude experienced during reinforcement learning in an experimental setting do not carry enough weight to make human learners negatively or positively biased across the board. After controlling for WoF (e.g. whether participants experienced win or a loss outcome on Day 1) and shape identity order (e.g. whether shape A would be encountered on a win or a loss day) and individual differences in how well participants learned the reward probability of the environment during the learning phase, there was no significant main effect of valence F(1, 64) = .307, p = .582, or reward probability F(5,320) = 1.542, p = .176) or WoF order/shape identity on participant choice behaviour (all p> .834, **Supplementary Figure 7A**). Within individual comparisons, we observed that participants were significantly positively biased for win shapes associated with 60% reward probability (t(73)=2.191, p=.03). Although our experimental design did not allow us to decompose this effect any further, it is important to highlight that these shapes were farthest away in proximity to the affective events (i.e. win and loss outcomes in the WoF draw) experienced during the learning/encoding stage, and were associated with highest level of expected uncertainty among all the better shapes. Further analysis of participant choice behaviour raising the possibility that expected uncertainty of the reward environment may drive non-linear preferences between affective memories is available in Supplementary Results.

### Human affective memories are represented non-linearly

Due to a high number of equal value comparisons reported in **Supplementary Table 1** and in **Supplementary Figure 7**, and also considering the inherent stochasticity in participant choice behaviour, it is difficult to establish a bird’s eye view on the organisation of human affective memories by solely relying on these comparisons. To be able to look beyond individual comparisons and construct a model of human value-based recall of affective memories which we probed with 400+ trials involving many random shape pairs (e.g. win 90% vs other day baseline 10%), we further analysed participant choice behaviour in the preference tests with computational modelling (see Supplementary Methods and Materials for details).

Here, it is important to highlight that in a large majority of the trials the expected value difference between the options were 0 (e.g. 60% win vs 60% loss shapes, **Supplementary Figure 1**), which would normally warrant random (i.e. 50-50) choices between these options, and consequently a benchmark log likelihood value of −.69 (i.e. log(.5)) for any decision model. First, we tested how well our stochastic choice model for the preference test which relies on the probability weighting function, performs against this benchmark. Across both

Study 1 and Study 2 this preference choice model performed significantly better than a random choice model (all t>8.2, all p<.001), meaning that the model can capture the subjective valuations underlying binary decision-making between affective memories. Our results demonstrate that when all trials and all possible comparisons between affective memories at different reward probability levels are concerned, discrete positive events (i.e. winning on a WoF draw) influence subsequent value-based recall of memories associated with the better option during RL (i.e. shapes associated with higher reward probabilities which were sampled more frequently during the encoding stage, based on difference in area under the curve Study 1 t(44)= 1.44 and 2.40, p=.15 and .02 (day 1 vs day 2 respectively); Study 2: t(68)=2.027, p=.047, Figure 3). This affective influence occurs in a manner that augments the subjective reward probabilities of these options during value-based recall (**Figure 3**). Although there were some differences in the execution of preference tests between 2 studies, we observed this positive induced bias consistently across 2 studies and 3 assessment time points.

**Figure 3.**
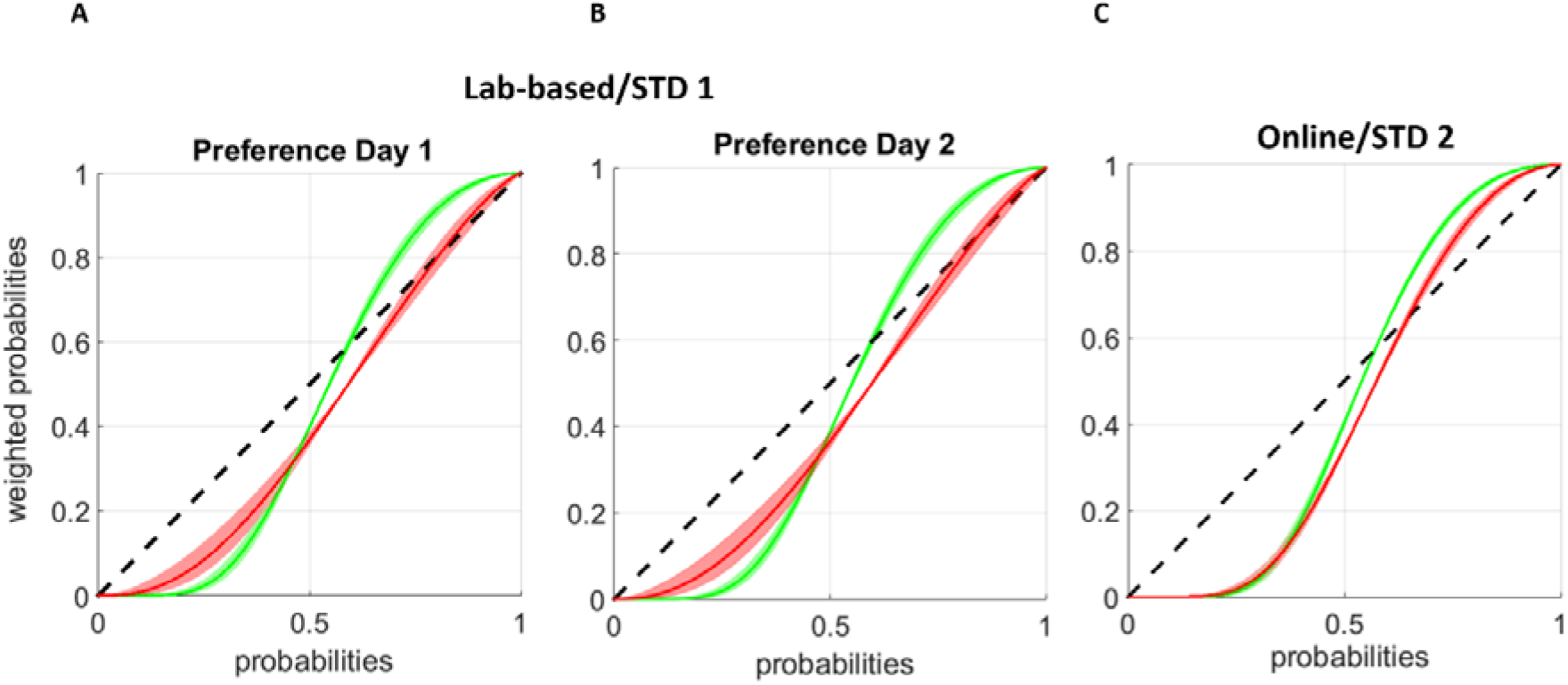
Probability weighting function demonstrating affective biases in memory-guided value-based decision-making. **(A-B)** In line with the model-free results reported in **Supplementary Figure 6**, model-based results show consistent effects between preference test days 1 and 2. The probability weighting function indicates that positive biases will have a stronger effect in contaminating neutral information associated with higher reward probability shapes (i.e. Baseline 70%), such that the reward probability associated with the baseline shape will be augmented when they are presented on the side associated with win shapes (green curve and SEM shading), whereas high probability baseline shapes will be under-weighted when they are presented on the side associated with loss shapes (red curve and SEM shading). Sidedness in stimuli presentation was counterbalanced across participants and between preference test days 1 and 2. This behaviour was reversed in the lower reward probability spectrum (i.e. for shapes associated with reward probability 30% and below). The trajectories of the weighting function capture the essence of all comparisons reported in Supplementary Table 1. **(C)** Perceived probabilities during value-based recall between affective memories in the online study (Study 2) in which all stimuli were presented randomly on each side of the screen. The difference between win and loss trajectories become more evident for higher probability shapes which were sampled more frequently during the encoding stage. Shading around the population mean denotes ± 1 SEM.

### Value-based decision-making between affective memories engages the pupil-linked central arousal systems

During the first preference test of Study 1, pupillometry data were collected across the entire decision process. We used a multiple linear regression model to quantify physiological response immediately before, during, and immediately after making choices between shapes learned following different WoF outcomes. Prior to choice, and even after controlling for the expected value difference between presented options as a proxy for choice difficulty, the expected value of chosen options estimated by the computational model reported above was significantly negatively correlated with pupil dilation (t(38) = −2.48, p = .018, **Figure 4A**). This means that choosing shapes associated with lower expected value leads to pupil dilation. After a choice had been made, affective memories had different physiological properties, and a rmANOVA indicated a significant timebin (i.e. every 1 second interval after decision-onset) by WoF outcome-valence interaction (F(9,333) = 2.28, p < .05, **Figure 4B**). This appeared to be driven primarily by the difference in peak pupil dilation between affective (i.e. win or loss) and blank shapes (a main effect of valence F(1,37)=3.865 and 5.997 (loss shapes versus win shapes respectively), p=.057 and .019). This neural response flips over from 2500 ms in the outcome delivery period (e.g. dilation to neutral and blank shapes increases over the average pupil dilation for loss shapes).

**Figure 4.**
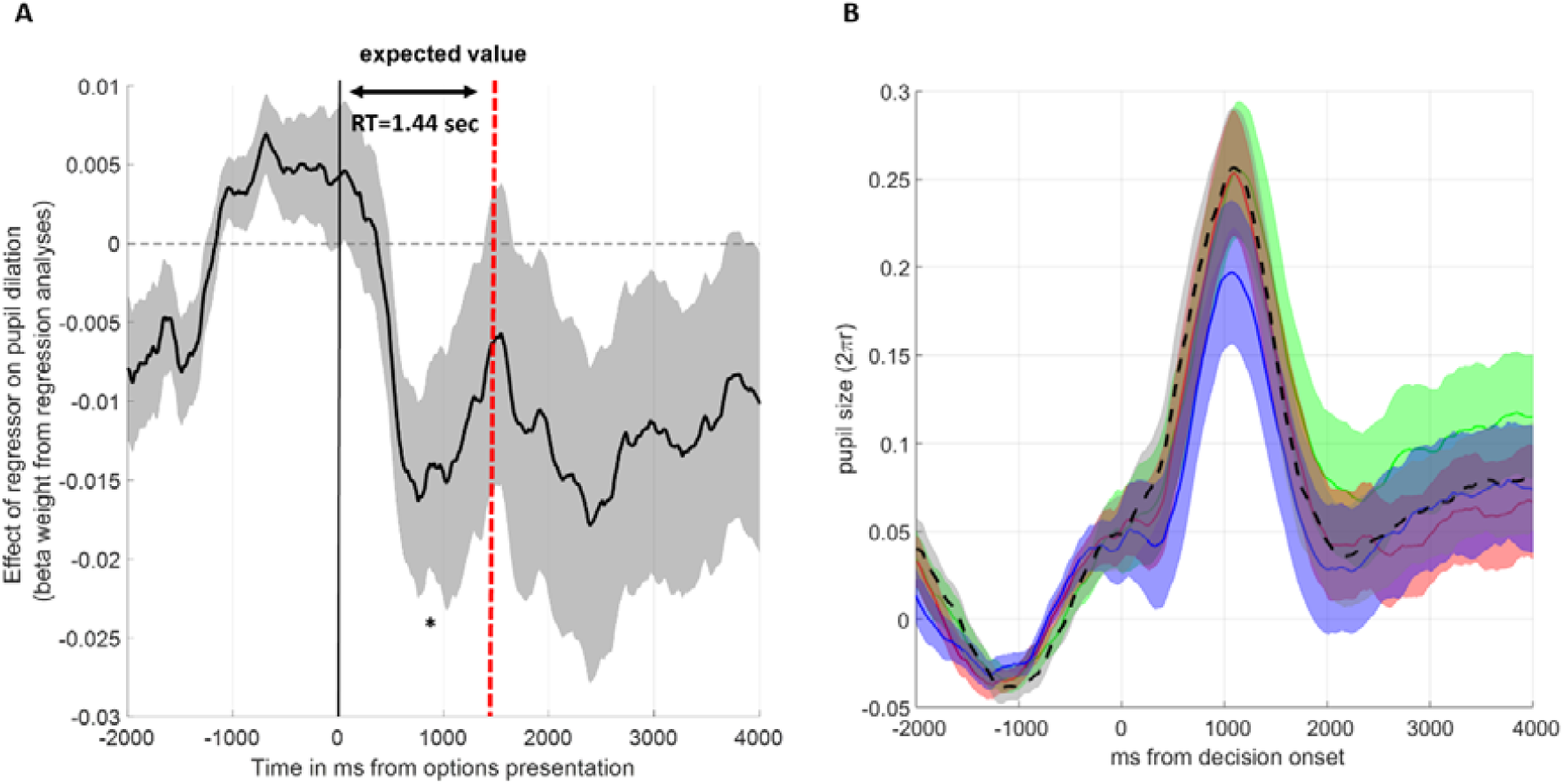
Pupillometry results. **(A)** A multiple linear regression analysis of the pupillary response suggests that pupil dilation negatively correlates with the expected values of chosen shapes based on the model shown in **Figure 4**. The vertical red dashed line marks the average response time (RT) during value-based decision-making. **(B)** During the outcome delivery period (i.e. once a decision has been made) affective memories lead to a larger pupil dilation relative to neutral memories (the difference between green/red lines versus the blue line). The significant valence x time bin interaction seems to arise from differential pupillary time courses between negative and neutral memories which cross over towards the end of the outcome delivery period. In both panels, shading around the population mean denotes ± 1 SEM. *p<.05.

## Discussion

In the current paper, we investigated mechanisms underlying value-based decision-making between affective memories formed under an RL protocol (**Figure 1**). Our findings from the preference tests suggest that human value-based decision-making between affective memories reveal stable preferences (**Supplementary Figure 6**). When only the pairwise comparisons of equal value options are concerned, discrete affective events of opposing valences do not carry enough weight to contaminate experimentally induced reward memories consistently across the reward probability spectrum (**Supplementary Table 1** and **Supplementary Figure 7**). However, when we also consider the global organisation of these affective memories (i.e. all cases where these memories were probed by randomly drawn options), our findings suggest that healthy volunteers retain positive biases for memories associated with better/higher probability options encoded through RL. We demonstrate that value-based decision-making between affective memories relies on nonlinear weighting of reward probabilities during recall (**Figure 3**). Taken together, these results illustrate that human memory-guided value-based decision-making is influenced by earlier experiences of discrete affective events and engages the pupil-linked central arousal systems prior to and after the decision onset (**Figure 4**).

In the current work, we investigated the degree to which nonclinical participants display a positive bias in value-based affective memory recall. Our reference point in designing this experiment was a rodent assay assessing the impact of rapid versus traditional antidepressants on a single negative memory relative to a single control condition^19^. However, in our experimental protocol, we probed a much larger pool of affective memories. For example, in the online study there were 24 abstract stimuli which could be uniquely paired with 23 other stimuli during the preference test, resulting in a total grid space of 552 combinations. When we consider this complexity and the global organisation of human affective memories, an overarching and conservative interpretation of our results is that nonclinical volunteers are overall positively biased in their value-based recall (**Figure 3**) and they maintain stable preferences between affective memories, at least during the first 48 hours following memory acquisition (**Supplementary Figure 6**). Although our approach captures the essence of the rodent assay, it reveals only the tip of the iceberg when it comes to fully understanding the organisation of experimentally induced affective memories in humans. There is substantial evidence, although limited to *deterministic* stimulus-outcome associations formed under conditioning, demonstrating that human learners store abstract knowledge in a grid-like code, in a manner that is similar to how the firing of entorhinal grid cells reflect spatial navigation in laboratory animals^27,28^. In our case, value-based recall demands navigating through an abstract reward probability space which is formed under a *stochastic* RL protocol and influenced by the valence of preceding affective events (i.e. WoF outcomes); it is therefore likely to have more uncertainty and nonlinearity in the way this information is stored. The second cognitive process relevant for understanding value-based recall is memory replay^29^. Previous research suggests that humans can simulate the timeline of events (e.g. remembering the loss on the WoF while recalling the reward probability associated with the better shape in the block immediately after) during memory recall and this can be detected through analysing the neural signature associated with different events happening in a sequence^30^. For example, a recent study demonstrated that events which generate large magnitude prediction errors create boundaries in memory formation^31^. In the context of our experimental protocol, the WoF draws were the affective events which arguably generated the largest magnitude of PEs and this might explain why we observed a nonlinearity in preferences for the some of the baseline shapes (**Supplementary Figure 7C-F**). Here, it is also worthwhile to note that our model-based analysis of individual RL blocks indicated that participants did not encode shape values through associations with their potential to generate large magnitude RPEs (i.e. Model 4), therefore it is more likely that in our experimental protocol event boundaries in memory emerged with respect to the WoF draw rather than learning individual reward associations within each block. Overall, these questions about grid-like organisation of human memory^32^ and memory recall through replay are timely topics within cognitive neuroscience and require further research, ideally using high-field MRI (further discussion available in Supplementary Methods and Materials).

Finally, our results demonstrate that value-based decision-making between affective memories engages the pupil-linked central arousal systems, with a negative correlation indicating that the pupil dilates more to chosen shapes with a lower expected value (**Figure 4A**). This is in line with recent computational work which showed that expected values of chosen options are negatively correlated with pupil dilation^33^. After the decision onset, the physiological response to affective memories are explained by a valence x time-bin interaction. Population averages of pupil traces for each outcome valence demonstrated that this significant interaction was driven by differential pupil dilation to negative versus neutral (i.e. blank WoF outcome) memories and between the early and late phase of the outcome delivery period (**Figure 4B**). Considering that pupil dilation is under the influence of a number of neurotransmitters such as norepinephrine, acetylcholine and serotonin^34^, our current work may be useful for understanding the effects of psychotropic compounds on affective memories. Although there is preliminary evidence to suggest that selective serotonin reuptake inhibitors induce a specific positive bias during value-based recall^22^, physiological correlates of this positive bias remain unknown. We think that future studies using imaging methods with high temporal resolution such as magnetoencephalography could be valuable in understanding neurotransmitter modulation of human memory systems.

## Funding and Disclosure

This work is funded by a joint grant from UK Medical Research Council and Janssen Pharmaceuticals awarded to CJH and SEM. CJH has received consultancy fees from P1vital Ltd., Janssen Pharmaceuticals, Sage Therapeutics, Pfizer, Zogenix and Lundbeck. SEM has received consultancy fees from P1vital, Zogenix, Sumitomo and Janssen Pharmaceuticals. EP has received consultancy fees from Janssen Pharmaceuticals. CJH and SEM hold grant income from UCB Pharma, Zogenix and Janssen Pharmaceuticals. CJH and SEM hold grant income from a collaborative research project with Pfizer. CG and HC do not declare any conflict of interest.

## Author Contributions

EP, CG, CJH and SEM designed the study. EP, CG and HC collected the data. EP analysed the data. All authors contributed to writing of the manuscript. Funders did not have any input in study design, analysis approach or decision to disseminate the results.

