## Supplementary Methods for "Value-based decision-making between affective memories"

### ***Supplementary Methods and Materials***

### ***Training phase***

In the lab-based study (Study 1) on each of the learning days, participants started with a baseline block (30 trials), followed by two more blocks (each 90 trials), with rest periods between the blocks. In the online study (Study 2), all blocks contained 40 trials each. The shapes in the RL tasks were selected from the Agathodaimon and Dingbat Cobogo fonts. During the RL blocks, participants were asked to learn the reward probabilities associated with each shape through trial and error. Participants were explicitly informed that on any given trial if one shape is rewarded the other one is not rewarded (such that the probabilities are p versus 1-p). A green frame appeared around the rewarded option to provide feedback on participant choice. For every correct choice made, participants collected 2 pence and they were asked to accumulate as much reward as possible. Participants started with £15 on day 1 and a running total at the bottom of the screen was updated at the start of each subsequent trial. In total, the participants were asked to learn reward probabilities associated with 14 shapes in the lab-based study and 24 shapes in the online study. Participant preferences between these shapes were later probed in the preference test which took place on the 4^th^ and 5^th^ days of the lab-based study and the 3^rd^/final day of the online study.

### ***Wheel of Fortune***

In order to influence the participants’ affective state, a single WoF draw was used in the break between the first and second blocks on each of the learning days. Participants were told that they could either win money, lose money, or receive nothing (blank). Unknown to the participants, the draw was not random but fixed, such that in the lab-based study each participant won (+£15), lost (-£11), or received a blank outcome (£0) across the 3 days. In the online study, we excluded the blank condition and focused only on win (+ £14) and loss (-£7) outcomes, and adjusted their outcome magnitudes considering the commonly observed salience differences between wins and losses^1^. To control for an effect of order, these outcomes were counterbalanced across participants. In Eldar and Niv ^2^ (2015), individual trials were rewarded with 25 cents and participants could either experience the loss or win of $7 after a single WoF draw (1/28 ratio). In contrast, we decided to increase the WoF outcome/reward ratio to ensure that these affective events (i.e. WoF outcomes) would feel more salient to participants. Therefore, in our experimental design, each correct prediction during RL led to a smaller reward (2p). As illustrated in **Figure 1A**, win and loss outcomes were sandwiched between large magnitudes of opposing outcomes, creating a near-miss effect aimed at strengthening the affective impact of the WoF.

### ***Happiness ratings***

During the learning days, participants were asked to report their current happiness. In the lab-based study (Study 1), we used a Likert scale from 1-9 (with higher numbers indicating greater happiness), whereas in the online study (Study 2) we used a visual analogue scale. Participants were asked to provide a happiness rating at different time points during the experiment, for example before starting the first RL block, more critically immediately before and after the WoF, and at the end of their training on any given day.

### ***Preference tests***

In the last phase of the experiment, we conducted preference tests. In order to assess the test-retest reliability of participant preferences we conducted the preference test twice, once on the 4^th^ day and again on the 5^th^ day in the lab-based study. Participants were presented with random pairs of all shapes they encountered during the training days and on each trial, were asked to choose the shape with the higher reward probability. The aim was to examine whether a WoF induced change of affective state would bias their valuations of the learned shapes. Written and spoken instructions were given and subjects were presented with a print-out of all the shapes (14 in Study 1 (lab) and 24 in Study 2 (online)) before the task began, to provide them with an opportunity to recall the learned shapes. On each day, preference tests consisted of 400/456 trials (lab-based versus online) presented across three blocks, and the order of these trials was randomised. The presented options prioritised randomly pairing equal probability shapes such that the value difference between the options would be near 0 (**Supplementary Figure 1**).

In the preference tests (both Study 1 and 2), participants did not receive any explicit feedback about whether their choices were correct or not, so they had to rely on what they had previously learned. In order to prevent further learning, participants' running total did not appear on the screen during a block of trials, but they did continue to accumulate money based on whether their choice was correct and the reward probability of the chosen shape. Their running total was only displayed between blocks to provide an indication of their performance in the previous block.

### ***Questionnaire measures***

In addition to the computerised task, participants were asked to respond to a series of self-report questionnaires on the first day of their testing, prior to completion of the first learning block. These questionnaires included: (1) Beck Depression Inventory (BDI-II)^3^, a standard measure of depression; (2) Spielberger State-Trait Anxiety Inventory (STAI)^4^, an anxiety measure comprising trait (anxiety proneness) and state (current state of anxiety) subscales; (3) Positive and Negative Affect Schedule (PANAS), assessing the feeling and expression of positive and negative emotions^5^; (4) Behavioural Activation/Behavioural Inhibition (BIS/BAS), reflecting aversive motivation and appetitive motivation^6^; (5) and the Mood Disorder Questionnaire (MDQ)^7^, a screening tool for Bipolar Disorder.

### ***Statistical analyses***

We used repeated-measures analysis of variance (rmANOVA) models to investigate the effects of our experimental manipulations. All significant main effects and interaction terms were followed up by post hoc tests, corrected for multiple comparisons using Bonferroni correction. All analyses were conducted on MATLAB and SPSS v25. In the lab-based study, 6 participants were excluded from the analyses of pupillometry data due to signal dropout affecting more than half of the trials, and in total 2 participants were excluded from the behavioural analysis (one person reported intentionally selecting the lower probability shapes during the training phase, the other person dropped out from the study before preference test 2). In total, 5 participants were excluded from the online study, 4 individuals who did not give more than 4 unique mood ratings across 16 assessment points, and one individual who consistently selected the lower probability shapes more frequently than the higher probability shapes. Across all statistical models, reward probability, valence and behaviour in pre-post-WoF blocks were entered as within subject factors, whereas WoF order (e.g. whether participants experienced win or a loss outcome on the 1^st^ day) and shape set which randomised the associations between shape identities and reward probabilities were entered as between subject factors.

### ***Modelling reinforcement learning***

Participants’ choice behaviour during the learning phase can be captured by a simple reinforcement learning model. For each participant, a learning rate (α) was estimated separately for each of the most rewarded shapes in each block of learning trials (e.g. a learning rate for the 90% reward probability shape which appeared immediately after losing on the WoF), using a Rescorla-Wagner learning rule which updates prediction errors (PEs) with a learning rate parameter^8^:

$p_{\left( i+1 \right)}= p_{(i)}+\alpha({Out}_{(i)}- p_{\left( i \right)})$	(1)

In this equation, $p_{(i)}$is the estimated reward probability associated with shape A, and *Out_(i)_* depicts whether shape A was rewarded or not on trial *i* (i.e. a vector of ones and zeroes where 1 indicates that the reward was associated with shape A). Considering that the reward probabilities were interdependent within each shape pair, the probability that shape B is rewarded on the current trial is 1-$p_{(i)}$. In Equation 1 $\alpha$is the learning rate estimated from participants’ choices.

We analysed participant choice behaviour with 4 variants of this model. All models had 3 free parameters in total, estimated individually from each participant’s choice behaviour.

Model 1 had two learning rates, updating participant’s belief about the probability of the better shape depending on the sign of the prediction error (i.e. learning rate for positive prediction errors and one learning rate for negative prediction errors).

Model 2 was based on a dynamic learning model recently reported as the best fitting model in another study investigating the effect of WoF outcomes on human learning and memory^9^:

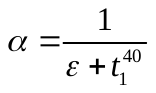
(2)

where ε>0 is a free parameter individually fitted to each participant data and additive trial numbers (*t*) in the denominator allow learning rates to diminish towards the end of the block, which is a widely accepted assumption for learning stable contingencies. The notation *t^40^* is because each RL block in the online study (Study 2) contained 40 trials. In this model, the WoF outcomes influence the subjective value of the rewards received during learning:

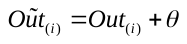
 (3)

where actual outcomes are transformed by an additive free parameter (θ), estimated between bounds [-2.5,2.5].

Model 3 had a single learning rate as shown in Eq.1. This model assumed that participants’ mood would influence RL on a trial-by-trial basis. Each participant’s mood trajectory within a block was retained by linearly interpolating between participant mood ratings which were captured by a visual analogue scale (VAS) at the beginning and the end each learning block (ratings as shown in **Supplementary Figure 2B).** The assumption tested by this model was that participants’ current mood would make it easier or more difficult to identify the better shape during the learning block. The effect of participant mood is implemented at the choice level, modulating decision values ($\Delta\tilde{v})$in a sigmoid function which generates choice probabilities for the better shape (*q_A_*):

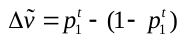
 (4)

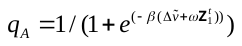
 (5)

where ω is a free parameter estimated in the same space as θ in Eq.3, and
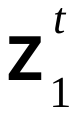
 is the vector of interpolated mood ratings, and β>0 is an inverse temperature parameter to capture the stochasticity in participant choices.

Finally, in Model 4 we tested whether separate learning rates assigned to update small and large magnitude of PEs can better account for participant choice behaviour during RL. A recent work demonstrated, within the context of Pavlovian conditioning, that experience of large PEs create event boundaries in human memory such that sequence of stimuli across a high PE event are recalled less accurately^10^. This model had the same 2 learning rate structure of Model 1, with an additional free parameter (0<κ<1) that serves as an individual sensitivity threshold to categorise trial-wise PEs into small versus large for each participant, and participant choice probabilities are generated by a sigmoid function driven by the value difference between the shapes (as in Eqs. 4 and 5).

In order to choose between these models we relied on the group-wise sum of Bayesian Information Criterion (BIC), which is a widely used metric for model selection^11^ (**Supplementary Figure 3)**.

### ***Modelling choice behaviour in the preference tests***

In this study, our primary research question is related to understanding value-based decision-making between affective memories, i.e. information encoded immediately after affective events such as winning or losing in a WoF draw. Considering that reward probabilities in RL blocks following the WoF draw were identical on each of the training days (e.g. 90% win probability in all RL blocks immediately after the WoF), a preference for one type of shape when participants are asked to choose between equal probability shapes can be understood in terms of probability weighting. This is because, when equal probability shapes are pitted against each other (e.g. a preference test trial in which participants are asked to choose between Win 90% versus Loss 90% shapes), preference for one type of shape means that either the reward probability associated with that shape is over-weighted, or the reward probability associated with the less preferred shape is under-weighted. This suggests that the 2-parameter probability weighting function^12^ can adequately capture participants’ preferences for shapes learned in different affective states. The strength of this approach is that the 2-parameter probability weighting function is free of any *a priori* assumptions about the shape of biases in probability weighting and can account for any non-linear weighting trajectories across the probability spectrum (i.e. between 0 and 1).

In the lab-based study (Study 1), we had abstract shapes falling into 4 different categories: baseline (i.e. pre-WoF shapes), win, loss, and blank depending on the outcome of the WoF on a given training day. This means that the ideal choice model guiding participants decisions can be expressed in terms of 8 parameters (2 parameters per shape category), accounting for probability weighting for each shape category. However, particularly in the baseline condition, our experimental design for the lab-based study did not allow enough coverage of the probability spectrum as the baseline condition only involved shapes with reward probabilities 70% versus 30%. These limitations, arising mainly from feasibility issues (i.e. limiting the number of shapes participants were asked to remember), meant that fitting a 2-parameter non-linear function to 2 data points on the probability spectrum (i.e. at 30% and 70%) is likely to give unreliable parameter estimates. To overcome this limitation, during preference testing we only presented win shapes on the left side and loss shapes on the right side, while blank and baseline shapes were presented on each side randomly. This side-specific stimuli presentation was counterbalanced across the participants and between preference test days 1 and 2. This approach allowed us to use the baseline shapes to increase the coverage of the probability spectrum at the data points which were otherwise missing in win and loss conditions, i.e. both conditions only had shapes associated with reward probabilities 10%, 20%, 80%, and 90%. Secondly, by presenting the blank outcome shapes randomly on each side we increased the number of trials with which we can assess the impact of affective events on low (i.e. 10% and 20%) and high (i.e. 80% and 90%) probability shapes. This approach allowed us to capture the degree to which negative and positive events contaminate otherwise neutral information (one widely accepted definition of affective bias) during value-based decision-making and to model participant choice behaviour in the preference test.

In the online study (Study 2), we did not have the blank/neutral condition and the reward probabilities associated with the shapes covered the probability spectrum more evenly (**Figure 1A**) which meant that we were able to present all shapes randomly on either side, overcoming some of the design limitations of Study 1.

We used the probability weighting function to estimate perceived probabilities associated with shapes presented on each side:

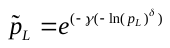
 (6)

Where p_L_ is the estimated probability of reward associated with any given shape, and it is based on participants’ learning behaviour linking the different stages of the experiment. The values of free parameters $\gamma$ and $\delta$ were estimated between 0 and 4.5, as setting up higher numbers for the upper boundary of these parameters has limited effect on the non-linear trajectories of the weighted probabilities. Then, the choice model assumes that participants will make decisions based on the expected value difference between available options presented on the left and the right side:

$\Delta\tilde{v}= \tilde{p}_{L}- \tilde{p}_{R}$	(7)

The trial-wise stochastic choice probabilities of each shape were generated using a sigmoid function^13^.

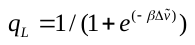
 (8)

where $\beta>0$ is the inverse temperature parameter, that governs the degree of stochasticity in participant choices. Values of $\beta$ tending towards 0 reflect an increase in stochasticity, whilst values of $\beta$ tending towards $\infty$ reflect more deterministic choices.

All free-parameters in computational models were estimated using a Bayesian model fitting procedure implemented in MATLAB. More specifically, the parameters were estimated by computing the full joint posterior probability of the models over the whole parameter space, and exact values of the free-parameters were computed by integrating the marginalised probability distributions of each parameter within specified parameter boundaries. Where applicable, learning rates were estimated in the inverse logistic space, probability weighting parameters were estimated in the normal space, and inverse temperature parameters were estimated in the log space. These are identical to parameter estimation procedures reported in our previous studies^14,15^.

### ***Pupillometry***

On the first day of preference tests (day 4) in the lab-based study, we collected pupillometry data to assess physiological response to affective memories during value-based decision-making. Participants' heads were placed at 70cm distance from the computer screen, stabilised by a chin rest. The eye-tracking system (Eyelink 1000 Plus; SR Research, Ottawa, Canada) was linked to the presentation computer through an ethernet connection. The sampling rate for pupillometry was set to 500 Hz recording from both eyes. The preference test was presented on a VGA monitor. A fixation cross marking the middle of the screen separated the presented pair of shapes.

Preprocessing of the pupillary data involved removing eye blinks that were identified using the built-in filter of the Eyelink system. A linear interpolation was implemented for all missing data points (including blinks). The resulting pupil trace was processed through a low pass Butterworth filter (cut-off of 3.75 Hz) and then z-transformed across the preference test session^16,17^. In order to assess phasic response to task-related variables, we performed baseline correction by subtracting the mean pupil size during the 2-second baseline period prior to our epochs of interest (i.e. decision and outcome) from each time point in the post-stimuli presentation period. Individual trials were excluded from the pupillometry analysis if more than 50% of the data from the outcome period had been interpolated^14,16^. The preprocessing resulted in a single set of pupil time-series per participant containing pupil dilation data for each of the included trials.

**Supplementary Results**

**Effect of Wheel of Fortune outcomes on happiness ratings**

In the lab-based study (Study 1), there was a significant main effect of WoF outcome on happiness (F(2,64) = 9.388, p < .001) and a significant WoF outcome by assessment time point interaction (i.e. the time point in which participants rated their momentarily happiness, F(2,64) = 33.467, p < .001). Importantly, there was no main effect of training order (e.g. whether participants experienced the win or the loss outcome on the WoF draw, F(5,32) = .964, p = .455) on any of the happiness ratings. As predicted, participants’ self-reported happiness ratings were significantly lower after losing the WoF draw (t = 5.59, p < .001, pairwise comparison of pre-WoF and post-WoF on loss day), whereas they were significantly higher on a WoF win day (t = -8.53, p < .001, pre-WoF vs. post-WoF, win day). These effects on happiness were maintained to the end of the final learning block on the loss day (t = 2.1, p < .05, comparing happiness pre-WoF to happiness at the end of the experiment on the loss day) but not the win day (t = -1.75, p = .088, comparing happiness pre-WoF to happiness at the end of the experiment on the win day), indicating that both winning and losing on the WoF induced significant changes in current happiness but that the effects of losing on the WoF had a longer-lasting impact on happiness than winning. The mood ratings assessed at the end of each day were significantly different between the conditions (F(2,134)=3.45, p=0.035). Pairwise comparisons indicated that happiness ratings were significantly lower at the end of the loss day relative to the win day (t(88)=2.7231, p=.008). Comparisons against the blank WoF were not significant (t(88)=1.5027, p=.137).

In the online study (Study 2), we mostly replicated these effects in a complementary rmANOVA model which suggested a significant main effect of WoF on happiness ratings (F(1, 67)= 5.431, p = .023), with a significant main effect of WoF outcome valence (i.e. win or a loss, F(1, 67)= 96.120, p<.001) and a significant valence by WoF interaction term (F(1, 67)= 133.155, p<.001). Again, there was no main effect of training order on the happiness ratings (F(1, 67)= 2.661, p = .108). Although participants mood ratings recovered from their lowest point which was immediately after a loss on the WoF, these still remained significantly lower on the loss day relative to the win day, even at the end of the experiment (t(136) = 3.675, p<.001, **Supplementary Figure 2B**).

**Model-based analysis of WoF effects on reinforcement learning**We modelled participant choice behaviour by fitting 4 different computational models (see Supplementary Methods and Materials for details). A well-established model selection method based on computing Bayesian Information Criterion values indicated that participants learned from rewarded and unrewarded outcomes (i.e. negative and positive prediction errors) with separate learning rates and made decisions between the shapes stochastically based on the expected value difference between the higher and lower probability shapes (i.e. Model 1, **Supplementary Figure 3**).

Fitting the same 2x3x2 rmANOVA model to positive learning rates (i.e. learning rate updating positive PEs) indicated a marginally significant main effect of WoF (F(1,65) = 3.904, p = .052),reflecting higher positive learning rates in the post-WoF blocks but no main effect of valence (F(1,65) = .117, p = .733, **Supplementary Figure 4A**). There were no significant interactions between these factors and no main effect of WoF order (F(1,65) = .094, p = .760). Analysing the negative learning rates in the same manner indicated a significant main effect of WoF (F(1,65) = 4.328, p = .041). There was also a significant main effect of outcome reward probability (F(1,65) =16.277 p<.001) and a marginally significant main effect of valence (F(1,65) = 3.068, p = . 085, **Supplementary Figure 4B**). Although there was no 3-way interaction, and no main effect of WoF order (F(1,65) = .422, p = .518), there was a significant interaction between outcome reward probability and valence (F(2,130)= 4.066, p = .019). Losing in the WoF reduced learning rates from negative prediction errors in blocks associated with higher expected uncertainty (i.e. reward probabilities 75% and 60%). Finally, repeating the same analysis for the inverse temperature term which governs choice stochasticity during RL indicated a significant main effect of WoF (F(1,65) = 7.503, p = .008) and a significant main effect of outcome reward probability (F(2, 130) = 75.499, p<.001), with no main effect of valence (F(1,65) = 2.162, p = .146, **Supplementary Figure 4C**). There were no significant interaction terms between these main effects and no main effect of WoF order (F(1,65) = 2.286, p = .135). Overall, inverse temperature estimates were higher in the post-WoF suggesting that participants were less stochastic in their choice behaviour. The results from a complementary model-free analysis based on participants’ choice switches is also reported in Supplementary Results.

**Model-free control analysis of WoF effects on reinforcement learning**

We conducted a model-free control analysis on the effects of WoF on human learning behaviour, focusing on participants’ probability of choice which following a “win” or a “no-win” outcome across different probability levels and pre-and post-WoF. In this analysis, probability of switching choice after a win outcome would be similar to the positive learning rates, whereas probability of switching choice after a no-win outcome would be similar to negative learning rates. Here is worthwhile to highlight that, this model-free analysis approach cannot capture the choice stochasticity effects, which are captured by the best fitting RL model. Inherent choice stochasticity commonly observed in human behaviour can confound the model-free analysis of choice switches.

We analysed participant choice behaviour by fitting a comparable 2x3x2 rmANOVA model to choice which probabilities following a win outcome, which indicated a significant main effect of WoF (F(1,65) = 6.018, p = .017) a significant main effect of valence (F(1,65) = 4.175, p = .045). There was no significant interaction between these factors (F(1,65) = 2.894, p = .094) and no main effect of WoF order (F(1,65) = .871, p = .354). In line with the model-based analysis indicating higher positive learning rates after winning in the WoF draw, there was a significant main effect of winning in the WoF on choice switch probabilities, which were consistently lower in the post-WoF blocks (F(1,65) = 7.488, p =.008, **Supplementary Figure 5A**).

Analysing the probability of choice which after a no-win outcome in the same manner indicated a significant main effect of WoF (F(1,65) = 12.649, p <.001). There was also no significant main effect of valence (F(1,65) = 2.288, p = .135). There was no significant interaction between these factors (F(1,65) = .613, p = .436) and no main effect of WoF order (F(1,65) = .661, p = .419). In line with the model-based analysis indicating lower negative learning rates after losing in the WoF draw, there was a significant main effect of losing in the WoF on choice switch probabilities, which were consistently lower in the post-WoF blocks (F(1,65) = 10.756, p =.002, **Supplementary Figure 5B**).

**Expected uncertainty of the reward environment may drive non-linear preferences between affective memories**

The second line of analysis was concerned with the direction of the influence of affective events created by the WoF draw. Our findings rule out the possibility that discrete affective events systematically contaminate memories which are encoded prior to them, as there were no consistent preferences between the baseline shapes learned before a win or a loss outcome on the WoF draw (F(1, 64) = .038, p = .846, **Supplementary Figure 7B**).

Analysing participant choice behaviour in favour of affective memories (i.e. post-WoF shapes) against baseline shapes both of the same and the opposing days in a 2x2x6 (valence x same versus other day x reward probability levels) rmANOVA indicated a significant main effect of reward probability (F(5, 320) = 12.311, p<.001) with no main effects of WoF and shape identity order, or any main effect of individual differences in learning (all p>.274). The main effect of reward probability was more evident for the high probability shapes, leading to non-linear preferences depending on the level of expected uncertainty in the learning environment. For example, irrespective of the WoF outcome valence, shapes learned immediately after the WoF draw (90% reward probability shapes) were preferred less relative to the equivalent baseline shapes, whereas those learned in the subsequent block (i.e. 75% reward probability shapes) were preferred consistently more than the equivalent baseline shapes (**Supplementary Figure 7C-F**).

**Supplementary Discussion**

One potential avenue which might improve our understanding of human memory formation and recall might be neuroimaging the habenular complex during RL and value-based decision-making between affective memories. A number of previous studies demonstrated that neurons in lateral habenula, an evolutionarily preserved subcortical structure located in the epithalamus, work in tandem with midbrain dopaminergic neurons during RL, particularly specialised in encoding the magnitude of negative PEs^31-33^. In the current work, we showed that when participants experience a large magnitude negative event prior to learning, they would be reluctant to update their beliefs when their reward expectations are violated in a novel learning environment (i.e. learning rates that update negative PEs, **Supplementary Figure 4B)**. A recent rodent study demonstrated that injecting ketamine into the lateral habenula attenuates the burst activity of the neurons in this region, rapidly ameliorating depressive symptoms^34^ in an animal model. Our experimental work and corresponding computational modelling suggest that discrete negative events influence components of human reinforcement learning which are shown to be encoded by lateral habenula neurons that can be modulated by NMDA receptor antagonists like ketamine. Therefore, a reverse translation of our experimental approach (i.e. using an instrumental conditioning framework) in rodents and recording the activity of the lateral habenula neurons could address some of the outstanding questions related to ketamine’s rapid antidepressant action such as its effects on affective information processing, and potentially memory formation.

**Supplementary Tables and Figures**

**Supplementary Table 1.** Equal value comparisons in the lab-based study (Supplement to SFig5)

| Day 1 | | | Day 2 | |
| --- | --- | --- | --- | --- |
| **Comparisons** | **t-statistic** | ***p*-value** | **t-statistic** | ***p*-value** |
| Win 90% vs Loss 90% | 0.36 | .72 | .51 | .61 |
| Win 10% vs Loss 10% | -.48 | .64 | -.31 | .76 |
| Win 80% vs Loss 80% | .62 | .54 | .57 | .57 |
| Win 20% vs Loss 20% | -2.95 | .005 | -5.45 | <.001* |
| Loss 90% vs Neutral 90% | 2.94 | .005 | 2.37 | .02 |
| Loss 10% vs Neutral 10% | -2.57 | .01 | -2.34 | .02 |
| Loss 80% vs Neutral 80% | .87 | .39 | 0.22 | .83 |
| Loss 20% vs Neutral 20% | 1.80 | .08 | 2.49 | .017 |
| Win 90% vs. Neutral 90% | 4.79 | < .001* | 4.19 | <.001* |
| Win 10% vs. Neutral 10% | -2.52 | .02 | -2.21 | .03 |
| Win 80% vs. Neutral 80% | 1.65 | .11 | .98 | .34 |
| Win 20% vs. Neutral 20% | -2.13 | .04 | -2.89 | .006 |

**df = 43, t-test pairwise comparisons, *.0042 (set level for 12 multiple comparisons Bonferroni corrected). Win 90% versus Neutral 90% is the only pairwise comparison which would survive Bonferroni correction over 2 preference tests.**

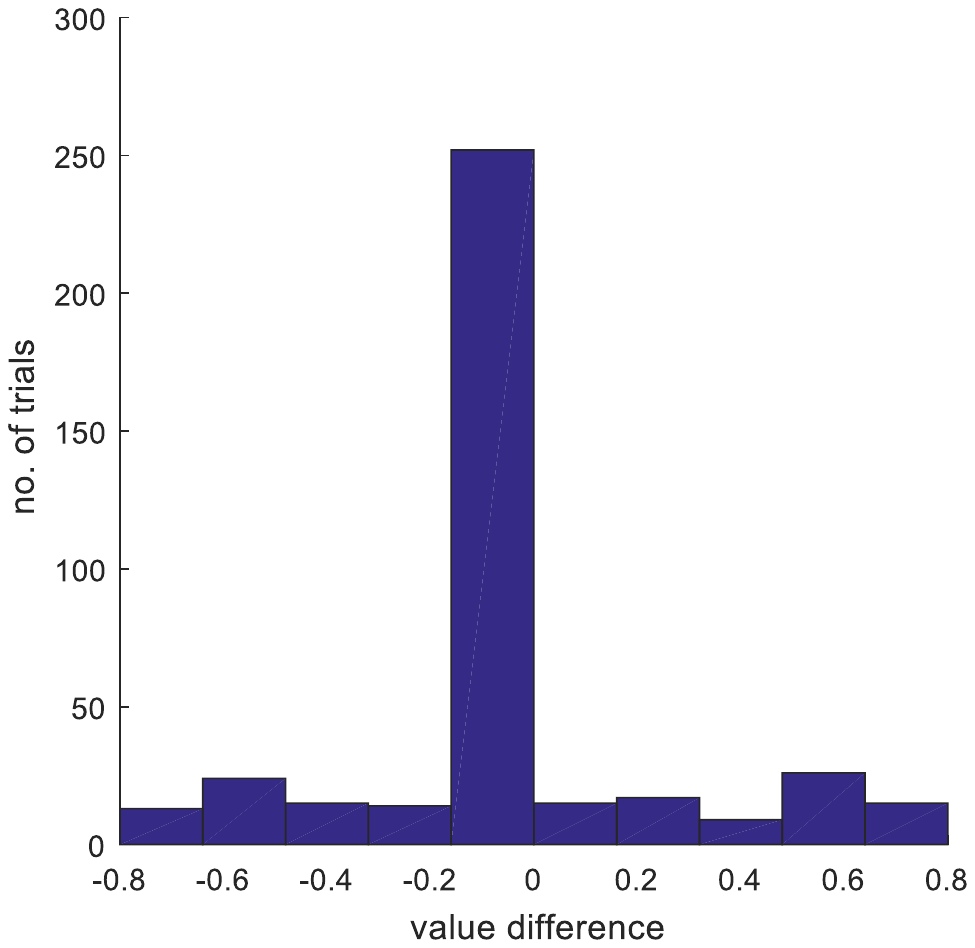

**Supplementary Figure 1**. **Value difference between shape options presented in the preference test.** Trials in the preference test were allocated to maximise the number of equal reward probability outcomes (i.e. where the value difference between presented shapes is zero).

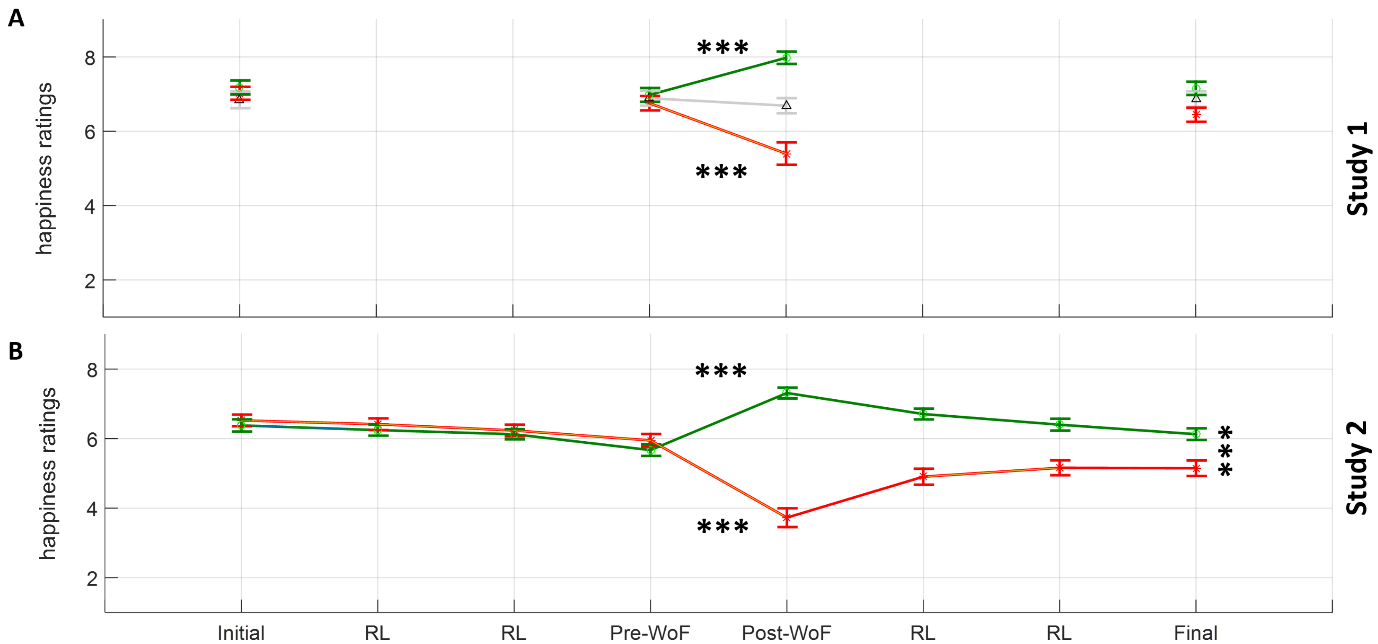

**Supplementary Figure 2**. **Happiness ratings** **(A)** In the lab-based study, participants’ current happiness significantly improved post-WoF on a win day (green line), while significantly worsening on a loss day (red line). On the day with a blank WoF draw (grey line), there was no significant difference between pre- and post-WoF happiness ratings. Moreover, initial and final happiness ratings were comparable on all but the loss day. On the day with the loss outcome, although current happiness recovered significantly from post-WoF, it was still significantly lower than participants’ initial rating. **(B)** These findings were replicated in the online study. The green line indicate mood ratings on the training day with a win on the WoF, whereas the red line indicate the mood ratings on the loss day. Across all panels, error bars reflect ±1 SEM. ***p<.001. RL blk: reinforcement learning block. In Study 2, the happiness ratings were made at the end of each block, apart from the initial rating which was made before participants started their daily training session.

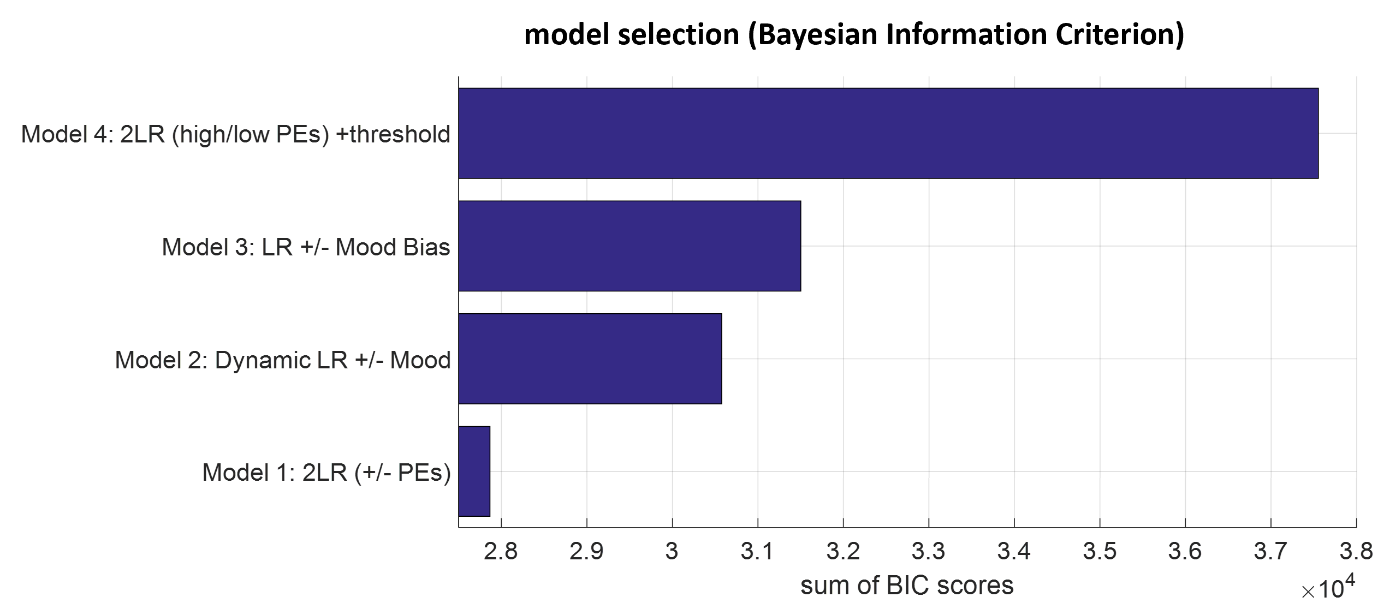

**Supplementary Figure 3**. Model selection based on group-wise sum of BIC scores comparing reinforcement learning models in the online study in which a model-free behavioural analysis based on participants’ frequency of choosing the better probability shape (in Figure 3B) suggested a significant main effect of WoF on human reinforcement learning. The best fitting model (Model 1) suggest that participants update their reward probability estimates by 2 distinct learning rates for negative and positive prediction errors.

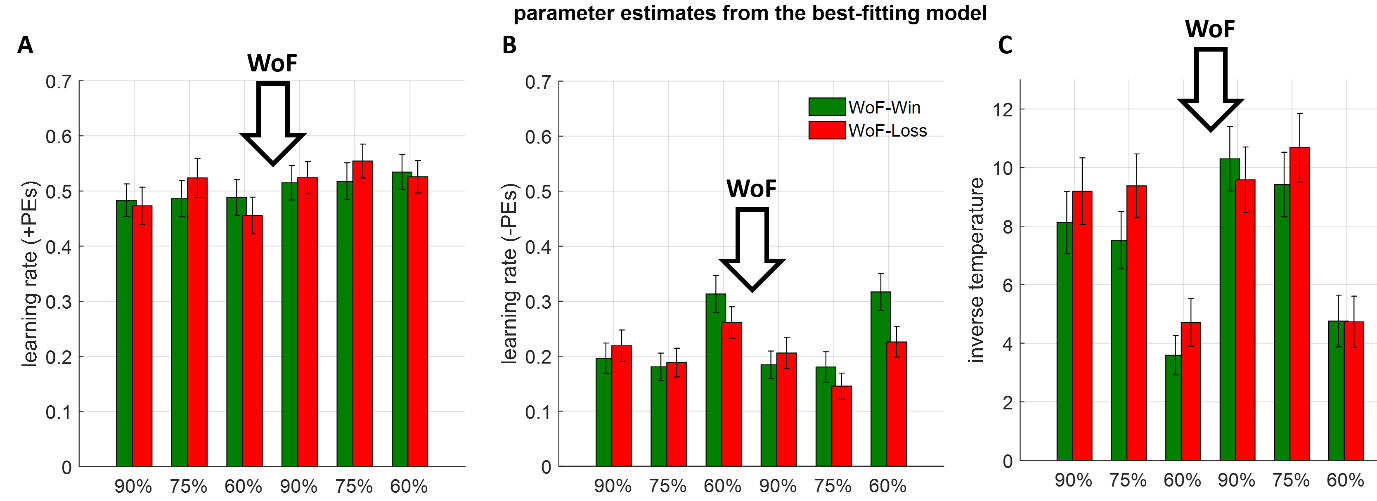

**Supplementary Figure 4**. **Summaries of parameter estimates from the best-fitting computational model** **(A)** Positive learning rates, by which participants update their beliefs about the reward probability following a reward outcome **(B)** Negative learning rates updating beliefs about the reward probability following a null/unrewarded outcome. **(C)** Inverse temperature term which governs choice stochasticity during reinforcement learning. Subsequent statistical analysis indicate that discrete affective events more strongly influence negative learning rates and choice stochasticity. Downward arrows with WoF indicate the point in which participants experienced the WoF draw within the course of their daily learning sessions. Across all panels, error bars reflect ±1 SEM. Results from a complementary model-free analysis focusing on participants probability of choice switches are reported as Supplementary Results and shown in Supplementary Figure 5.’

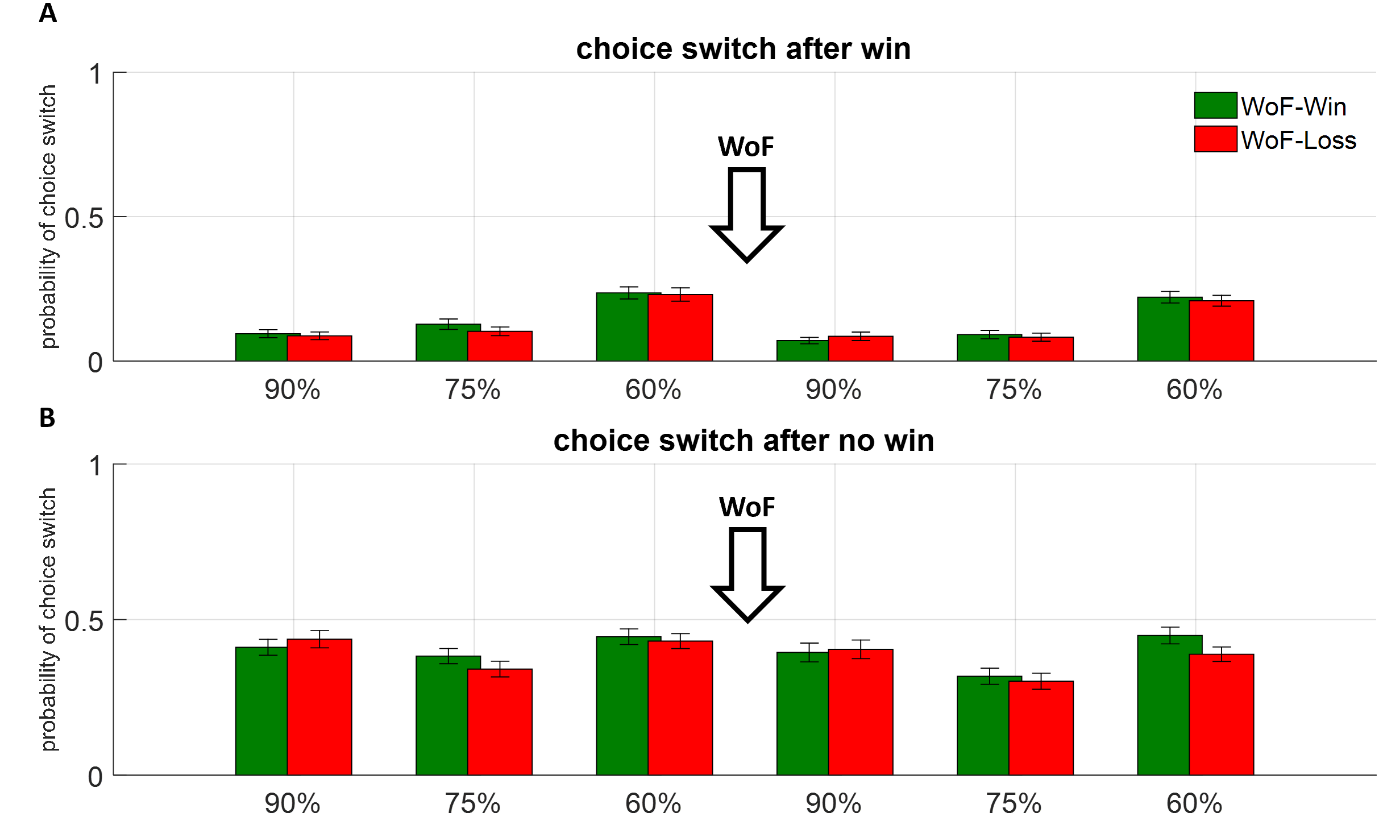

**Supplementary Figure 5**. **Model-free analysis of participant learning behaviour** **(A)** Probability of choice which following a win outcome. **(B)** Probability of choice which following a no-win outcome. Downward arrows with WoF indicate the point in which participants experienced the WoF draw within the course of their daily learning sessions. Across both panels, error bars reflect ±1 SEM.

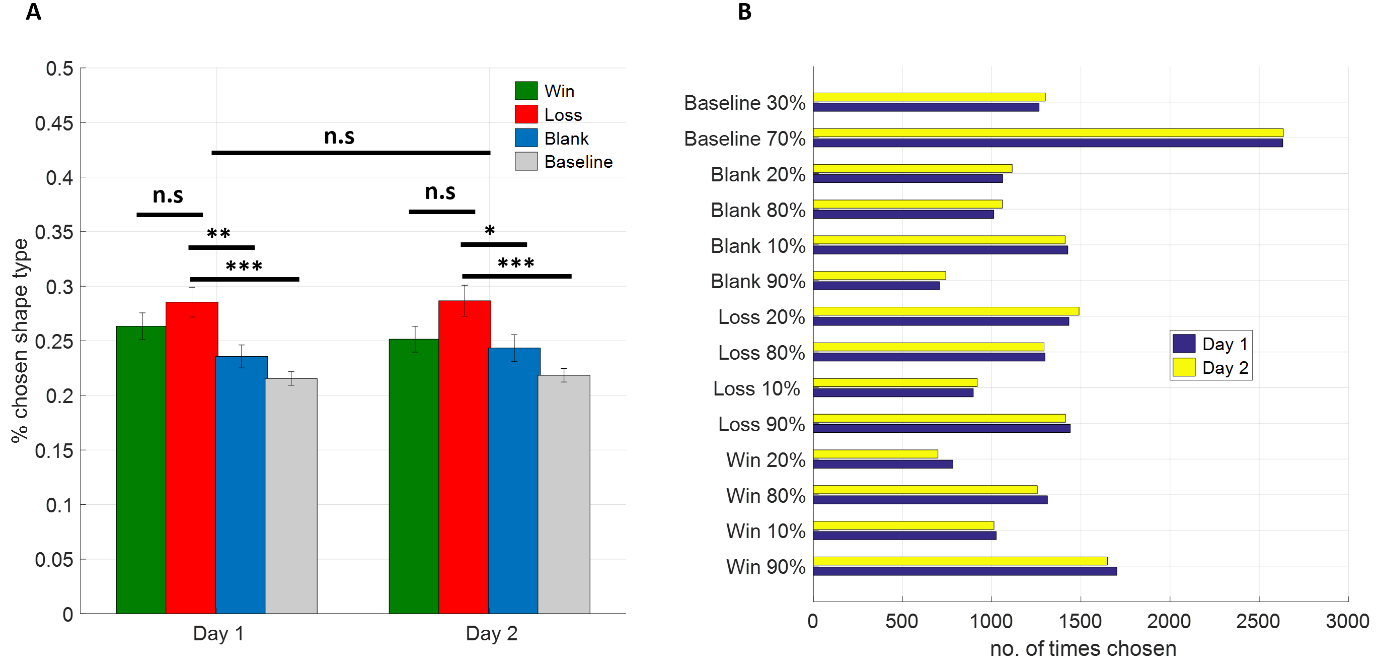

**Supplementary Figure 6. Chosen shape types in the lab-based study**. **(A)** Overall summary of chosen shapes between day 1 and day 2, sorted by WoF valence. Loss shapes were selected more frequently than other shapes, whereas there was no significant difference in participant choice behaviour between day 1 and day 2. Error bars denote ±1 SEM. **(B)** Detailed summary of how frequently individual shapes were chosen. The increase in preference for loss shapes reported in panel A appears to be driven by a stronger preference for Loss 20% shapes relative to Win 20% shapes. ***p<.001, **p<.01, *p<.05, n.s: not significant.

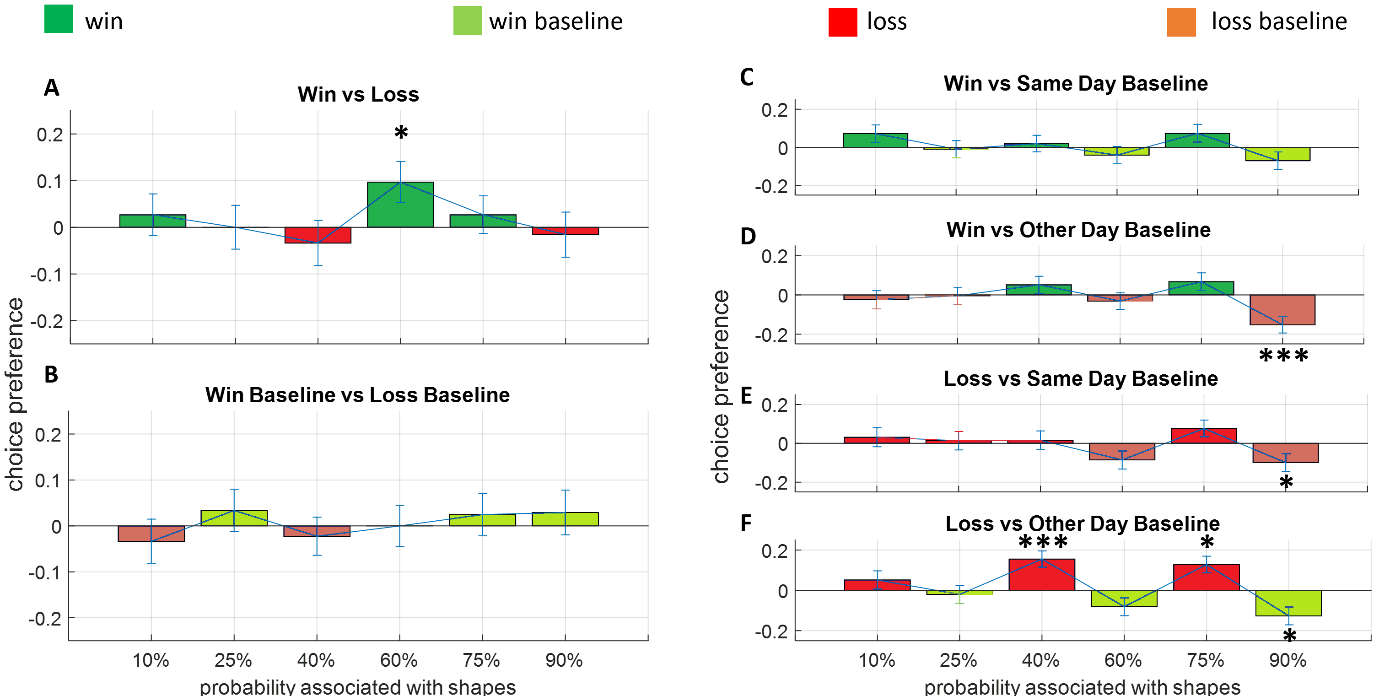

**Supplementary Figure 7. Comparisons between equal value shapes. (A)** Shapes learned after a win or a loss WoF draw**. (B)** Shapes learned prior to a win or a loss WoF draw. **(C-F)** Shapes learned before versus after a win or a loss WoF draw including same and other day baseline comparisons. Values across all panels are normalised, meaning that positive values on the y-axis indicate a preference for affective (i.e. post-WoF) shapes. Across all panels, error bars reflect ±1 SEM. ***p<.001, *p<.05, uncorrected. All reward probabilities are reordered in linearly increasing order. Bar graphic face colours follow the legend above top panels.
